# Inflammatory signaling drives SQSTM1-dependent suppression of interferon-γ–induced MHC class I surface expression

**DOI:** 10.64898/2026.10.23.761340

**Authors:** Sho Yamashita, Hiroe Shuto, Aiko Hirayama, Atsushi Shimauchi, Takahiro Utsumi, Toshifumi Ninomiya, Kazuya Tsubouchi, Kohei Otsubo, Daisuke Shibahara, Eiji Iwama, Isamu Okamoto, Kentaro Tanaka

**Affiliations:** Department of Respiratory Medicine, Graduate School of Medical Sciences, Kyushu University, Fukuoka, Japan; Department of Respiratory Medicine, Fukuoka Red Cross Hospital, Fukuoka, Japan

## Abstract

An immune-inflamed tumor microenvironment is characterized by the coexistence of interferon-γ (IFN-γ) and inflammatory cytokines, yet how these opposing signals are integrated by cancer cells to regulate tumor immunogenicity remains poorly understood. Here, we investigated interleukin-1β (IL-1β) on IFN-γ–induced major histocompatibility complex class I (MHC-I) expression in non–small cell lung cancer (NSCLC) cells using molecular and cell biological approaches. IL-1β suppressed IFN-γ–induced surface MHC-I expression despite preserved or enhanced expression of antigen presentation–related genes, indicating post-transcriptional regulation. This effect emerged 12–24 h after stimulation and was accompanied by accelerated MHC-I turnover without altering total protein abundance. RNA sequencing identified selective induction of autophagy-related genes specifically under combined IFN-γ and IL-1β stimulation. Immunofluorescence and biochemical analyses demonstrated intracellular redistribution of MHC-I to autophagosome–lysosome compartments. Mechanistically, the selective autophagy receptor SQSTM1 associated with MHC-I and was required for its surface downregulation, whereas depletion of NBR1 or inhibition of autophagy or lysosomal function failed to restore MHC-I expression, indicating a nonredundant role for SQSTM1. These findings identify post-translational MHC-I trafficking as a previously underappreciated regulatory layer linking inflammatory signaling to the canonical IFN-γ–MHC-I axis and reveal an SQSTM1-dependent mechanism controlling MHC-I surface availability under immune-inflamed tumor conditions.

## Introduction

The introduction of immunotherapeutic agents, such as programmed cell death–1 (PD-1) pathway inhibitors, has greatly improved the prognosis of individuals with many types of malignant tumors, including non–small cell lung cancer (NSCLC) ^1^ ^2^ [^3^ ^4,5^. Further advances in immunotherapy are expected to include the development of novel agents such as bispecific antibodies and chimeric antigen receptor–T cell (CAR-T) therapy ^6^.

Such advances will require optimization of the induction of cytotoxic CD8^+^ T cells (CTLs), essential effector cells of antitumor immunity, with a focus on elucidation of the molecular immune mechanisms involved in carcinogenesis and the development of resistance to immunotherapy ^7^. Defects in interferon-γ (IFN-γ) signaling, neoantigen depletion, various immune checkpoints, and tumor-mediated immune suppression are representative targets of such research. Decreased expression of major histocompatibility complex class I (MHC-I) molecules is thought to be one of the most important factors associated with immunosuppression leading to cancer development. Antigen presentation by MHC-I determines the specificity of CD8^+^ T cells and is essential for their activation and proliferation ^8^. Professional antigen-presenting cells such as dendritic cells as well as certain cancer cells are capable of processing self- or tumor antigens to form stable MHC-I–peptide complexes at the cell surface that engage CD8^+^ T cells through antigen-specific T cell receptors and thereby trigger CTL responses. Mutations or loss of expression of essential genes whose products contribute to various stages of MHC-I antigen presentation—such as MHC-I and β2-microglobulin ^9^ ^10^ ^11^, transporters associated with antigen processing (TAP1 and TAP2) ^12^ ^13^, calreticulin and TAPBP (which mediate peptide transport and loading) ^14^ ^15^ ^16^, CITA (also known as NLRC5) and IRF2 (MHC-I gene transcriptional activators) ^17^ ^18^, and the kinases JAK1 and JAK2 (which mediate MHC-I induction) ^19^—have been implicated in tumor progression as well as the development of resistance to immune checkpoint blockade ^7^ ^20^. Genetic alterations do not fully represent the mechanisms that lead to low immunogenicity of tumors, however, with additional components associated with dysregulation of MHC-I antigen presentation in the complex context of the tumor microenvironment (TME) likely to play an important role.

The immune landscape of the TME varies among tumor types ^21^. In NSCLC, which manifests a high level of immune cell infiltration (an immune-inflamed TME), various cytokines are produced by immune cells surrounding the tumor cells. Among these cytokines, IFN-γ produced by T cells plays a central role in tumor immune surveillance by increasing the expression of MHC-I molecules in tumor cells and thereby activating the antigen presentation machinery. Increased surface expression of MHC-I promotes efficient recognition of tumor cells by CTLs and consequent antitumor immune responses ^11^, with loss of MHC-I expression constituting a central mechanism of tumor immune evasion as a result in part of escape from CD8⁺ T cell–mediated killing. However, whether inflammatory cytokines in addition to IFN-γ contribute to such regulation, and whether tumor cell–intrinsic mechanisms govern IFN-γ–mediated MHC-I expression, have remained poorly understood.

We previously performed a comprehensive analysis of cytokines produced in the TME of NSCLC and found that the inflammatory cytokine interleukin-1β (IL-1β) is abundant and contributes with IFN-γ to the synergistic induction of programmed cell death–ligand 1 (PD-L1) in tumor cells ^22^. IL-1β was found to be produced mostly by tumor-associated macrophages (TAMs), which participate in the establishment of an immunosuppressive TME that promotes tumor progression ^23^ ^24^ ^25^. We therefore hypothesized that IL-1β might play a key role in regulation of MHC-I expression at the cell surface, and we here evaluated the effect of IL-1β on IFN-γ–induced MHC-I expression in lung cancer cells. Multiple analyses revealed that IL-1β suppresses the IFN-γ–induced expression of MHC-I at the tumor cell surface. Unexpectedly, we found that this suppressive effect was not attributable to changes in gene transcription or total protein abundance of MHC-I, but rather to posttranslational regulation. Comprehensive RNA–sequencing (seq) analysis, subcellular localization, and loss-of-function approaches revealed that this effect of IL-1β is mediated by promotion of autophagy-lysosome trafficking of MHC-I that is dependent on the selective autophagy receptor SQSTM1.

## Materials and Methods

### Cell culture and reagents

H1437, H2122, HCC827, and H1650 cells were obtained from American Type Culture Collection, and II-18 cells from the RIKEN BRC cell bank. These cell lines were cultured in RPMI 1640 medium (Gibco) supplemented with 10% fetal bovine serum and 1% penicillin-streptomycin (Gibco). All cells were maintained under a humidified atmosphere of 5% CO_2_ at 37°C. Recombinant human IL-1β (R&D Systems, #201-LB) and recombinant human IFN-γ (PeproTech, #300-02) were dissolved in phosphate-buffered saline (PBS) containing 0.1% bovine serum albumin. Chlorpromazine (#S5749), ZCL278 (#S7293), methyl-β-cyclodextrin (#S6827) and 3-Methyladenine (#S2767) were obtained from Selleck, and chloroquine (#54-05-7) from Tokyo Chemical Industry. Each of these inhibitors was dissolved in dimethyl sulfoxide (DMSO).

### Immunoblot analysis

Cells were washed with ice-cold PBS and then lysed in RIPA buffer (Thermo Fisher Scientific) containing protease and phosphatase inhibitor cocktails for preparation of whole cell lysates. A cytoplasmic extract was prepared with the use of NE-PER Nuclear and Cytoplasmic Extraction Reagents (Thermo Fisher Scientific, #78833), and cell surface proteins were isolated with a Pierce Cell Surface Biotinylation and Isolation Kit (Thermo Fisher Scientific, #A44390). All samples were fractionated by SDS-polyacrylamide gel electrophoresis on a 10% gel, and the separated proteins were transferred to a polyvinylidene difluoride membrane. The membrane was incubated overnight at 4°C with primary antibodies to MHC-I (#88274), to β-actin (#4970), to E-cadherin (#3195), to TAP1 (#49671), to TAP2 (#25657), to Tapasin (#66382), to LMP2 (#87667), to LMP7 (#13635), to NBR1 (#20145), to SQSTM1 (#88588), or to LC3B (#3868) all of which were obtained from Cell Signaling Technology, as well as with those to SUSD6 (#HPA030464) obtained from Sigma-Aldrich. The membrane was then incubated for 1 h at room temperature with horseradish peroxidase–conjugated secondary antibodies to rabbit (Cytiva, #NA9340) or mouse (Cytiva, #NA931VS), after which immune complexes were detected with the use of Pierce ECL Plus Western Blotting Substrate (Thermo Fisher Scientific) or a SuperSignal West Dura Extended Duration Substrate Kit (Thermo Fisher Scientific). Blot images were captured with a ChemiDoc XRS+ system (Bio-Rad).

### RT-qPCR analysis

Total RNA was extracted from cells with the use of an RNeasy Mini Kit (Qiagen, #74904), and portions of the RNA (100 ng) were subjected to reverse transcription (RT) with PrimeScript RT Master Mix (Takara Bio, #RR036A). The resulting cDNA was subjected to quantitative polymerase chain reaction (qPCR) analysis with the use of TB Green Premix Ex Taq II (Takara, #RR820A) and a Thermal Cycler Dice Real Time System (Bio-Rad). The qPCR primers (forward and reverse, respectively) included those for 18S rRNA (5΄-ACTCAACACGGGAAACCTCA-3΄ and 5΄-AACCAGACAAATCGCTCCAC-3΄), MHC-I (5΄-AGTGGGCTACGTGGACGACA-3΄ and 5΄-ATGTAATCCTTGCCGTCGTA-3΄), TAP1 (5΄-GCAGTCAACTCCTGGACCACTA-3΄ and 5΄-CAAGGTTCCCACTGCTTACAGC-3΄), TAP2 (5΄-ATGCCCTTCACAATAGCAGCGG-3΄ and 5΄-CCAAAACTGCGAACGGTCTGCA-3΄), ERAP1 (5΄-GTCTGTCAGTGTGACCCATCCT-3΄ and 5΄-CTGAGCAGGATTTTCCACAGGTG-3΄), ERAP2 (5΄-CTGTGACCTGAACCATGCTCCT-3΄ and 5΄-TCCATCCTGCTGTTGTCTGAGC-3΄), TAPBP (Tapasin) (5΄-GAGCCTGTTCTCATCACCATGG-3΄ and 5΄-GTAGGCAAAGCTCAAGTCCAGC-3΄), ERp57 (5΄-GTCAGCCACTTGAAGAAGCAGG-3΄ and 5΄-TAGGAACTCGGAGTGAGCCTCA-3΄), and SUSD6 (5΄-AGTGGAAACCAGCCATGGAG-3΄ and 5΄-CCACTATAGACAGCGTGGGG-3΄). The amount of each mRNA was quantitated by the ΔCt method and normalized by that of 18S rRNA.

### Flow cytometric analysis

The surface expression of MHC-I in NSCLC cells was quantified by flow cytometry. For surface staining, cells were washed with FACS buffer (PBS containing 2% fetal bovine serum), incubated for 1 h at 4°C with fluorophore-conjugated antibodies diluted to the appropriate concentration in FACS buffer, and washed again twice with FACS buffer. For intracellular staining, performed after fixed in 4% paraformaldehyde (Nacalai Tesque, #09154-56) for 15 min at room temperature, cells were permeabilized with 0.3% Triton X-100 (Nacalai Tesque, #35501-02) for 10 min at 4°C, and incubated for 1 h at 4°C with antibodies diluted in FACS buffer. Cells were stained with antibodies to HLA-A,B,C (BioLegend, #311409) or an allophycocyanin-conjugated mouse IgG2a isotype control (BioLegend, #400220). All samples were analyzed by flow cytometry with a FACSLyric instrument (BD Biosciences), and the data were processed with Flowjo (v10.10.0) software. The mean fluorescence intensity (MFI) ratio was calculated as the MFI for MHC-I divided by that for the isotype control.

### Immunofluorescence analysis

Cells were seeded on 12-mm-diameter cover glasses (Matsunami) in a 24-well plate, fixed in 4% paraformaldehyde (Nacalai Tesque, #09154-56) for 15 min at room temperature, and permeabilized (or not) with 0.3% Triton X-100 (Nacalai Tesque, #35501-02) for 10 min at 4°C. They were washed with ice-cold PBS, incubated for 1 h at room temperature with blocking buffer (KAC, UKB40), and exposed overnight at 4°C to primary antibodies in a humidified chamber (Incubation Chamber, Cosmo Bio, #10DO). The cells were again washed with ice-cold PBS before incubation for 1 h at room temperature with fluorescent dye–conjugated secondary antibodies. Nuclei were counterstained with 4′,6-diamidino-2-phenylindole (DAPI) (Nacalai Tesque, #12745-74). The primary antibodies included those to MHC-I (Cell Signaling Technology, #88274; 1:250 dilution), to calnexin (Proteintech, #10427-2-AP; 1:400), to GM130 (Cell Signaling Technology, #12480; 1:2000), to TGN46 (Proteintech, #13573-1-AP; 1:2000), to EEA1 (Cell Signaling Technology, #3288; 1:100), to LAMP1 (Cell Signaling Technology, #9091; 1:100), to E-cadherin (R&D Systems, #AF748; 1:20), and to LC3B (Cell Signaling Technology, #3868; 1:2000). Secondary antibodies included those to mouse (Abcam, #ab150114; 1:500), rabbit (Abcam, #ab150077; 1:500), or goat (Abcam, #ab150129; 1:200) IgG.

### Microscopy

All images were acquired with an all-in-one fluorescence microscope (BZ X800, Keyence) equipped with a Plan Apochromat ×100 objective (numerical aperture of 1.45; BZ-PA100, Keyence). Optical sectioning was applied to capture cross-sectional images of specimens at the focused depth so as to avoid blurred, out-of-focus images. Optical-sectioning images of cells were obtained by *z*-stacking from the top to the bottom of each cell at 0.5-μm intervals with the use of a sectioning module (BZ-H3XF, Keyence).

### Immunofluorescence staining and quantification of cell surface MHC-I

Cells were fixed with 4% paraformaldehyde without permeabilization and stained with anti-MHC-I antibody. The expression level of cell surface MHC-I was quantified as the average fluorescence intensity per cell using a BZ-X Analyzer software (Keyence).

### Immunofluorescence staining and quantification of intracellular MHC-I

Cells were fixed and permeabilized, followed by co-staining for MHC-I and E-cadherin. Using the E-cadherin signal as an indicator of the cell membrane, the region was eroded inward by 5 pixels to exclude the cell surface using a BZ-X Analyzer software (Keyence). The resulting region was defined as the cytoplasmic compartment, and intracellular MHC-I expression was quantified by measuring the mean fluorescence intensity within this region on a per-field basis using a BZ-X Analyzer software (Keyence).

### Cycloheximide (CHX) chase assay

H1437 cells were seeded at a density sufficient to ensure ∼80% confluency after 72 h. At 24 h after seeding, the cells were incubated for 24 h with IFN-γ in the absence or presence of IL-1β and then for an additional 12 or 24 h in the additional presence of CHX (Sigma-Aldrich, #239765) at 50 µg/ml. The cells were then harvested for preparation of protein lysates and immunoblot analysis.

### Transfection of siRNAs

H1437 cells were seeded in six-well plates and subjected to transient transfection for 24 h with small interfering RNAs (siRNAs) mixed with the RNAiMAX reagent (Thermo Fisher Scientific). The siRNAs included those specific for LC3B, NBR1, SUSD6, or SQSTM1 (Thermo Fisher Scientific) as well as a negative control siRNA (Japan Bio Services). The cells were then treated with IFN-γ in the absence of presence of IL-1β for 48 h, harvested, and subjected to flow cytometry.

### Immunoprecipitation

Cells were lysed in Pierce IP Lysis Buffer (Thermo Fisher Scientific, #87787) containing protease and phosphatase inhibitors (Nacalai Tesque), the lysates were centrifuged at 13,000 × *g* for 10 min at 4°C, and the resulting supernatants were subjected to immunoprecipitation overnight at 4°C with constant mixing, with the addition of primary antibodies to MHC-I (CST, #88274) or to immunoglobulin G (IgG) (CST, #5415S). The antibody-protein complexes were then incubated with Pierce Protein A/G Magnetic Beads (Thermo Fisher Scientific, #88802) for 1 h at room temperature with mixing. The bead-bound complexes were washed three times with Tris-buffered saline with the use of a magnetic column. Immunoprecipitated proteins were eluted from the beads by mixing for 10 min with Pierce IgG Elution Buffer, pH 2.0 (Thermo Fisher Scientific, #21028), and the eluate was neutralized with Neutralization Buffer (1 M Tris-HCl, pH 8.5; Nippon Gene, #316–90405) and subjected to immunoblot analysis.

### Kyoto Encyclopedia of Genes and Genomes (KEGG) pathway enrichment analysis

Differentially expressed genes (DEGs) identified from bulk RNA sequencing were subjected to KEGG pathway enrichment analysis using Metascape (https://metascape.org), a web-based gene annotation and analysis tool. Pathways with a P value < 0.05 were considered significantly enriched.

### Statistical analysis

Statistical analysis was performed with GraphPad Prism software (RRID: SCR_002798). All in vitro experiments were conducted at least three times. Comparisons among more than two groups were performed by one-way analysis of variance (ANOVA) followed by Šídák’s multiple-comparison test or Dunnett’s test, as appropriate. A *p* value of <0.05 was considered statistically significant.

## Results

### IL-1β inhibits IFN-γ–induced MHC-I expression at the surface of lung adenocarcinoma cells

We first examined the effects of IL-1β and IFN-γ on the surface expression of MHC-I in NSCLC cells by flow cytometry. Whereas IFN-γ induced a significant increase in such expression in H1437, II-18, H2122, and HCC827 cells (Fig. 1A). Although IL-1β alone did not significantly alter surface MHC-I levels compared to untreated controls (data not shown), IL-1β significantly attenuated the stimulatory effect of IFN-γ on surface MHC-I expression in each cell line. Immunofluorescence staining similarly revealed that IFN-γ alone increased the intensity of MHC-I signals that overlapped with the cell membrane marker E-cadherin in H1437 and II-18 cells, and that this effect of IFN-γ was inhibited by IL-1β (Fig. 1B, C). Immunoblot analysis showed that, whereas IL-1β did not substantially affect the total abundance of MHC-I protein in IFN-γ–stimulated H1437 and II-18 cells, it suppressed the IFN-γ–induced increase in the amount of MHC-I in a surface protein fraction of these cells (Fig. 1D), consistent with the findings of flow cytometry and immunofluorescence analysis. In contrast, in the NSCLC cell line H1650, IL-1β did not attenuate the IFN-γ–induced increase in surface MHC-I expression as detected by flow cytometry (Supplementary Fig. S1A) or immunoblot analysis (Supplementary Fig. S1B). RT-qPCR analysis showed that the amount of MHC-I mRNA was increased by IFN-γ stimulation in H1437, II-18, and H1650 cells, and that this effect was not influenced by the presence of IL-1β (Fig. 1E, Supplementary Fig. S1C). Together, these results revealed that IFN-γ–induced MHC-I expression at the cell surface was attenuated by IL-1β in certain NSCLC cells, and that this inhibitory effect of IL-1β was not mediated at the level of transcription or translation.

**Figure 1.**
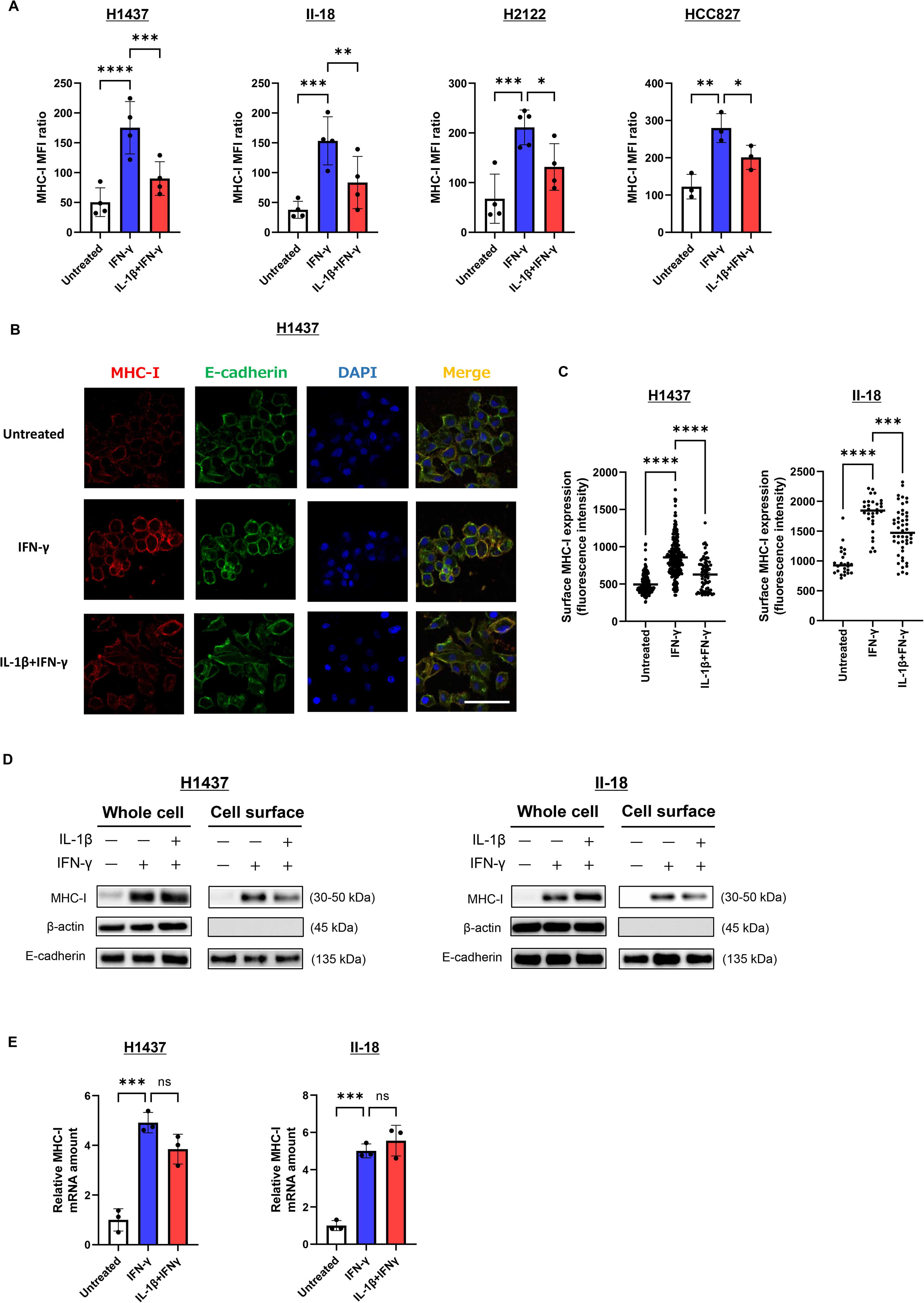
MHC-I expression enhanced by IFN-γ stimulation is suppressed by combined IL-1β stimulation in Lung Adenocarcinoma. IFN-γ–induced MHC-I expression at the cell surface is suppressed by IL-1β in lung adenocarcinoma cells. **A:** Flow cytometric analysis of surface MHC-I expression in H1437, II-18, H2122, and HCC827 cells treated with IFN-γ (50 ng/ml) either alone or together with IL-1β (50 ng/ml) for 48 h. **B:** Representative immunofluorescence analysis of MHC-I and E-cadherin in H1437 cells fixed after treatment as in **A**. Nuclei were stained with DAPI. Scale bar, 50 μm. **C:** Quantification of the fluorescence intensity of surface MHC-I expression in H1437 or II-18 cells from images similar to those in **B**. Data are shown for individual cells examined in three biological replicates, with the horizontal bars indicating median values. **D:** Immunoblot analysis of whole cell lysates and a cell surface protein extract prepared from H1437 or II-18 cells treated as in **A**. β-actin and E-cadherin were examined as cytoplasmic and cell surface marker proteins. Data are representative of three biological replicates. **E:** RT-qPCR analysis of MHC-I mRNA in H1437 or II-18 cells treated as in **A**. Quantitative data in **A** and **E** are means ± SD from three or four independent experiments. \**p* < 0.05, \*\**p* < 0.01, \*\*\**p* < 0.001, \*\*\*\**p* < 0.0001; ns, not significant (one-way ANOVA followed by Dunnett’s test).

### Inhibition of IFN-γ–induced surface MHC-I expression by IL-1β is independent of the canonical antigen-processing machinery and ER-Golgi transport

To investigate whether targeting of canonical mechanisms of antigen presentation might account for the downregulation of surface MHC-I expression by IL-1β in IFN-γ–treated cells, we examined the expression of proteins involved in MHC-I antigen presentation, including the proteasome subunits LMP2 and LMP7 (which cleave ubiquitinated antigens into peptides), TAP1 and TAP2 (which transport cleaved peptides into the endoplasmic reticulum [ER]), ERAP1 and ERAP2 (which further process peptides within the ER), and TAPBP (Tapasin) and ERp57 (which facilitate peptide loading onto MHC-I). RT-qPCR and immunoblot analyses revealed that the expression of these various molecules in H1437 cells was similar in the presence of IFN-γ either alone or together with IL-1β, with the exception that the IFN-γ–induced increase in the amount of ERAP2 mRNA was significantly enhanced by IL-1β (Fig. 2). In II-18 cells, whereas the expression of the various proteins was largely unaffected by IL-1β in the presence of IFN-γ, the abundance of the corresponding mRNAs was further increased by IL-1β (Fig. 2), possibly reflecting a mechanism to compensate for the reduced surface expression of MHC-I.

**Figure 2.**
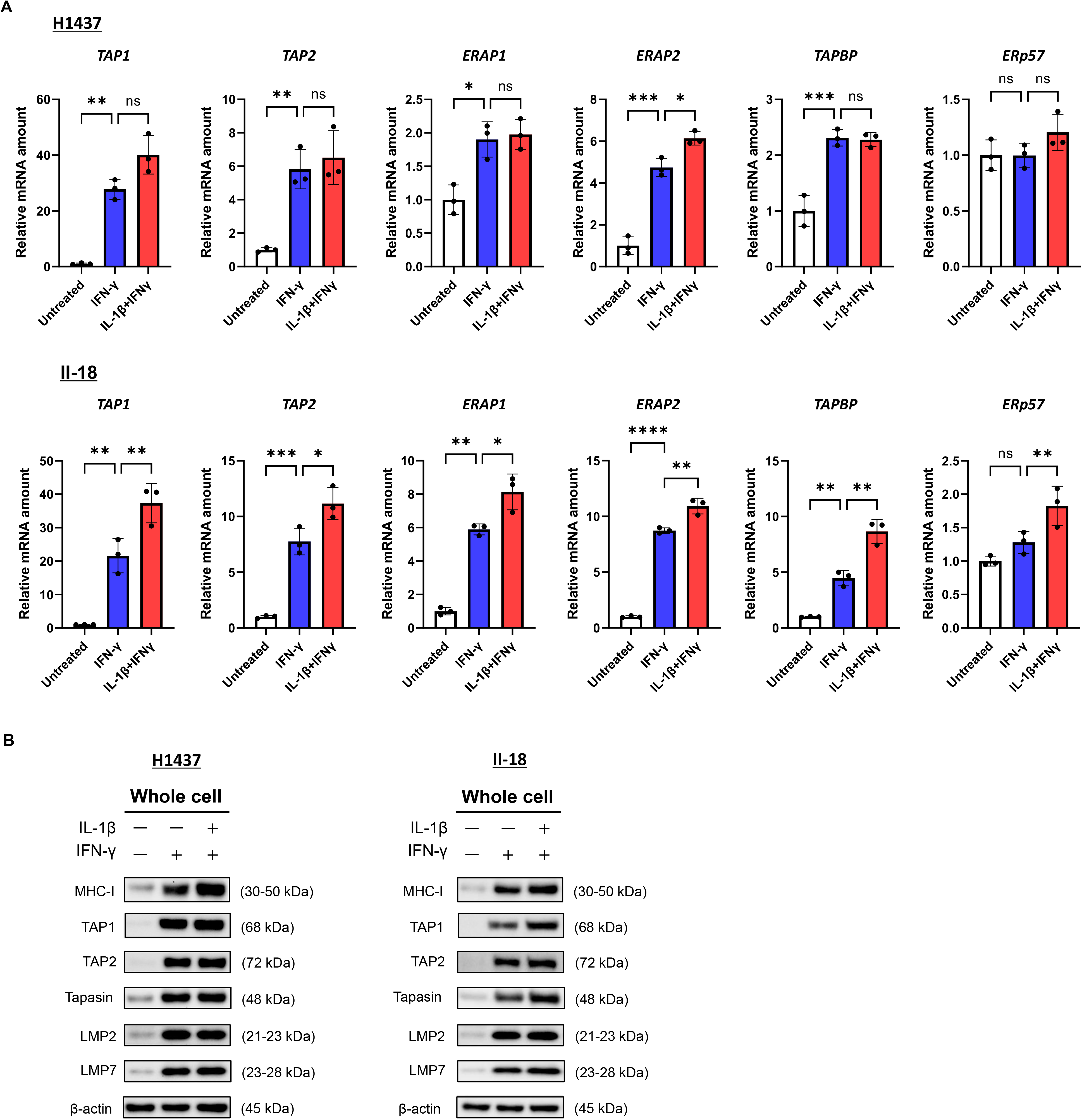
Genes’ expression related with canonical antigen-presenting mechanisms was maintained under IL-1β and IFN-γ co-stimulation. Expression of molecules related to canonical antigen presentation mechanisms is maintained in IFN-γ–treated cells in the presence of IL-1β. H1437 or II-18 cells incubated in the absence or presence of IFN-γ (50 ng/ml) and IL-1β (50 ng/ml) for 48 h were subjected to RT-qPCR analysis of the indicated mRNAs (**A**) as well as to immunoblot analysis of the indicated proteins (**B**). The immunoblots are representative of three biological replicates. Data in **A** are means ± SD from three independent experiments. \**p* < 0.05, \*\**p* < 0.01, \*\*\**p* < 0.001, \*\*\*\**p* < 0.0001; ns, not significant (one-way ANOVA followed by Dunnett’s test).

We next examined whether cytokine stimulation might differentially affect the transport of MHC-I from the ER through the Golgi apparatus to the cell membrane. Immunofluorescence staining revealed that the colocalization of MHC-I with the ER marker calnexin did not differ among untreated H1437 cells and those treated with IFN-γ alone or together with IL-1β (Supplementary Fig. S2A, B). In contrast, colocalization of MHC-I with the cis-Golgi marker GM130 or the trans-Golgi marker TGN46 was increased in IFN-γ–stimulated cells compared with control cells, and this effect of IFN-γ on MHC-I colocalization with TGN46 was significantly attenuated by IL-1β (Supplementary Fig. S2C–F). Collectively, these results suggested that the suppression of IFN-γ–induced MHC-I surface expression by IL-1β was not attributable to retention in the ER or Golgi apparatus or to inhibition of membrane transport. Instead, consistent with the observed effects on the expression of genes related to antigen presentation, membrane transport mechanisms may actually be enhanced by IL-1β in the presence of IFN-γ.

### IL-1β induces time-dependent intracellular accumulation of MHC-I in IFN-γ–stimulated cells

On the basis of our results thus far, we speculated that MHC-I might accumulate within NSCLC cells treated with both IFN-γ and IL-1β. To address this possibility, we assessed MHC-I expression by flow cytometry, immunofluorescence staining, and immunoblot analysis. Simultaneous flow cytometric analysis of surface and intracellular MHC-I expression revealed that, whereas IL-1β attenuated the IFN-γ–induced increase in MHC-I expression at the surface of H1437 cells, it enhanced the IFN-γ–induced increase in intracellular MHC-I expression (Fig. 3A). The ratio of intracellular to surface MHC-I expression was thus significantly increased in the presence of IL-1β. Consistent with these results, immunofluorescence analysis revealed that cytoplasmic MHC-I signals were more pronounced in cells treated with both IFN-γ and IL-1β than in those treated with IFN-γ alone (Fig. 3B, C). In addition, immunoblot analysis of a cytoplasmic fraction showed that, whereas the upregulation of MHC-I by IFN-γ was not affected by IL-1β at 24 h, it was enhanced at 48 h (Fig. 3D). These data thus indicated that MHC-I was internalized in response to IL-1β treatment in IFN-γ–stimulated cells, and that the extent of this internalization increased over time.

**Figure 3.**
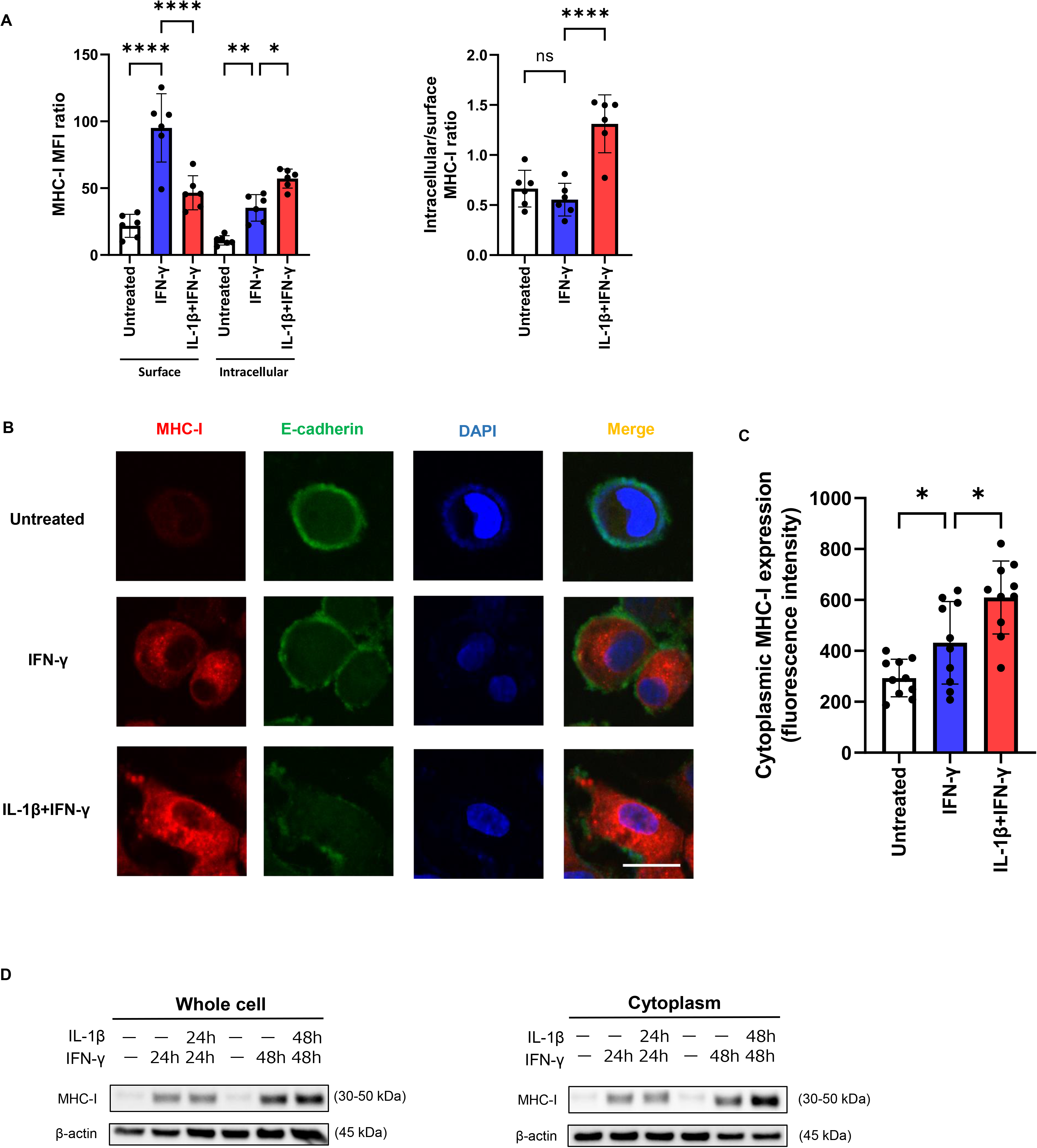
The intracellular localization of MHC-I increases in a time-dependent manner with combined IL-1β stimulation. IL-1β increases the intracellular localization of MHC-I in IFN-γ–treated cells in a time-dependent manner. **A:** Flow cytometric analysis of surface and intracellular MHC-I in H1437 cells treated with IFN-γ (50 ng/ml) either alone or together with IL-1β (50 ng/ml) for 48 h. Data are means ± SD from six independent experiments. **B:** H1437 cells treated as in **A** were fixed and permeabilized before immunofluorescence staining of MHC-I and E-cadherin. Nuclei were stained with DAPI. Scale bar, 10 μm. **C:** Quantification of cytoplasmic fluorescence intensity for MHC-I in images as in **B**. Data are means ± SD for individual cells in three biological replicates. **D:** Immunoblot analysis of whole cell lysates and a cytoplasmic protein extract prepared from H1437 cells incubated in the absence or presence of IFN-γ (50 ng/ml) and IL-1β (50 ng/ml) for 24 or 48 h. The immunoblots are representative of three biological replicates. \**p* < 0.05, \*\**p* < 0.01, \*\*\*\**p* < 0.0001; ns, not significant (one-way ANOVA followed by Dunnett’s test).

### IL-1β promotes endosome-to-lysosome trafficking and lysosomal degradation of MHC-I independently of canonical endocytic pathways

To investigate the fate of internalized MHC-I molecules, we examined their subcellular localization by immunofluorescence analysis of the early endosome marker EEA1 (early endosome antigen 1) and the lysosome marker LAMP1 (lysosomal-associated membrane protein 1) in cells exposed to the lysosome inhibitor chloroquine. Compared with IFN-γ stimulation alone, costimulation with IL-1β and IFN-γ markedly increased the colocalization of MHC-I with both early endosomes and lysosomes in H1437 cells (Fig. 4A–D), indicative of enhanced internalization of MHC-I and a consequent reduction in its surface expression. This pattern of MHC-I localization as well as its intracellular accumulation over time (Fig. 3D) suggested that costimulation accelerates lysosomal degradation of MHC-I. Treatment of cells with the protein synthesis inhibitor CHX at 24 h after the onset of cytokine stimulation revealed that the abundance of MHC-I at 12 h was similar in cells treated with IFN-γ either alone or together with IL-1β. In contrast, at 24 h, the amount of MHC-I was markedly attenuated by costimulation with IFN-γ and IL-1β (Fig. 4E). These findings indicated that the increased colocalization of MHC-I with lysosomes under the condition of costimulation results in its enhanced degradation and reduced overall protein stability.

**Figure 4.**
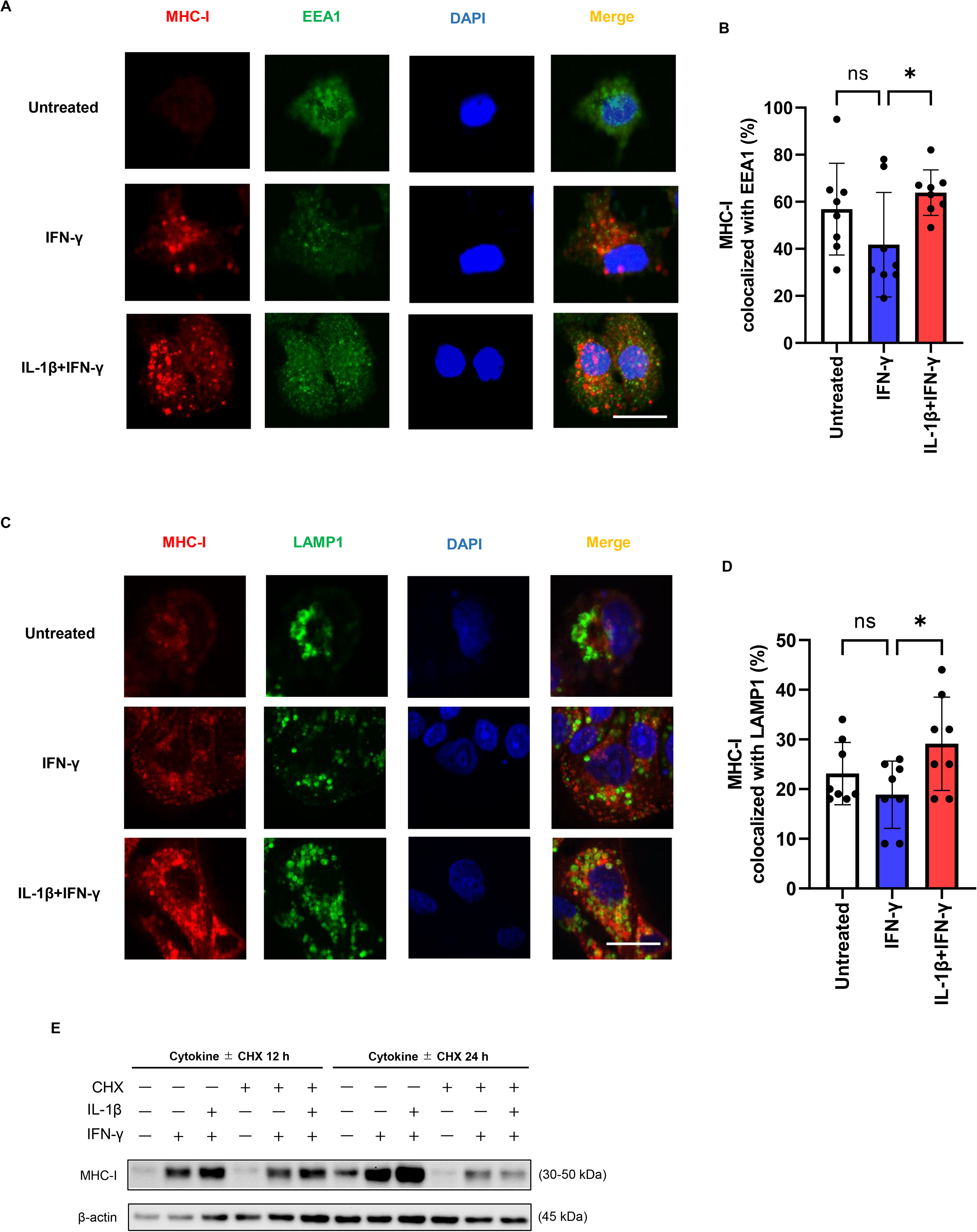
IL-1β accelerates lysosome-dependent degradation of MHC-I following intracellular internalization. IL-1β and IFN-γ costimulation promotes MHC-I association with autophagy-related organelles. **A, C:** H1437 cells incubated with or without IFN-γ (50 ng/ml) and IL-1β (50 ng/ml) and in the presence of the lysosome inhibitor chloroquine (50 μM) for 48 h were fixed, permeabilized, and subjected to immunofluorescence analysis of MHC-I and the early endosome marker EEA1 (**A**) or the lysosome marker LAMP1 (**C**). Nuclei were stained with DAPI. Scale bars, 10 μm. **B, D:** Quantification of the percentage colocalization of MHC-I with EEA1 (**B**) or LAMP1 (**D**) for images as in **A** and **C**, respectively. Data are means ± SD for individual cells from three independent experiments. \**p* < 0.05; ns, not significant (one-way ANOVA followed by Dunnett’s test). **E:** H1437 cells were treated with IFN-γ (50 ng/ml) alone or together with IL-1β (50 ng/ml) for 24 h and then incubated in the additional absence or presence of CHX (50 µg/ml) for 12 or 24 h. Whole cell lysates were then subjected to immunoblot analysis of MHC-I. The immunoblots are representative of three biological replicates.

Cell migration–inducing and hyaluronan-binding protein (CEMIP) was previously shown to promote MHC-I internalization via clathrin-dependent endocytosis ^26^. Endocytosis is mediated by multiple pathways including caveola-dependent, lipid raft-dependent and cdc42-dependent mechanisms in addition to the clathrin-dependent mechanism. To examine whether these endocytic routes are associated with MHC-I internalization in cells exposed to both IL-1β and IFN-γ, we investigated the effects of pharmacological inhibitors targeting these pathways. H1437 cells were stimulated with cytokines and subsequently treated with chlorpromazine (clathrin- and caveola-dependent endocytosis inhibitor), methyl-β-cyclodextrin (caveola- and lipid raft–dependent endocytosis inhibitor), or ZCL278 (cdc42 inhibitor). The inhibitory effect of IL-1β on surface MHC-I expression induced by IFN-γ was unaffected by chlorpromazine or ZCL278 and was actually enhanced by methyl-β-cyclodextrin, likely as a result of a nonspecific cytotoxic effect (Supplementary Fig. S3). These findings thus indicated that canonical endocytosis pathways are not necessary for the internalization of MHC-I induced by IL-1β in IFN-γ–treated cells.

### IL-1β and IFN-γ costimulation induces autophagy activation and MHC-I association with autophagic structures

To elucidate the molecular mechanisms underlying the internalization of MHC-I in cells exposed to IFN-γ and IL-1β, we performed bulk RNA-seq analysis of H1437 cells treated with both cytokines and those treated with IFN-γ alone. In addition to many pathways related to IL-1β and IFN-γ stimulation and signaling, the autophagy pathway was significantly upregulated in the cells exposed to both cytokines (Fig. 5A). Genes whose expression was increased under the costimulation condition included canonical autophagy-related genes such as *ATG7* and *MAP1LC3B* (encoding LC3B, a key component of autophagosome membranes) as well as *NBR1*, *SUSD6* (sushi domain containing 6), and *SQSTM1* (sequestosome 1, or p62) (Fig. 5B). Consistent with these findings with H1437 cells, these autophagy-associated genes also showed increased expression in II-18 cells treated with both IL-1β and IFN-γ (GEO database). Immunofluorescence analysis further revealed that costimulation increased autophagy activity—as evidenced by increased LC3B signal intensity—and increased the extent of MHC-I colocalization with autophagic structures, compared with IFN-γ treatment alone, in H1437 cells (Fig. 5C). These results suggested that IL-1β promotes MHC-I internalization and likely degradation via activation of the autophagy machinery in IFN-γ–treated cells.

**Figure 5.**
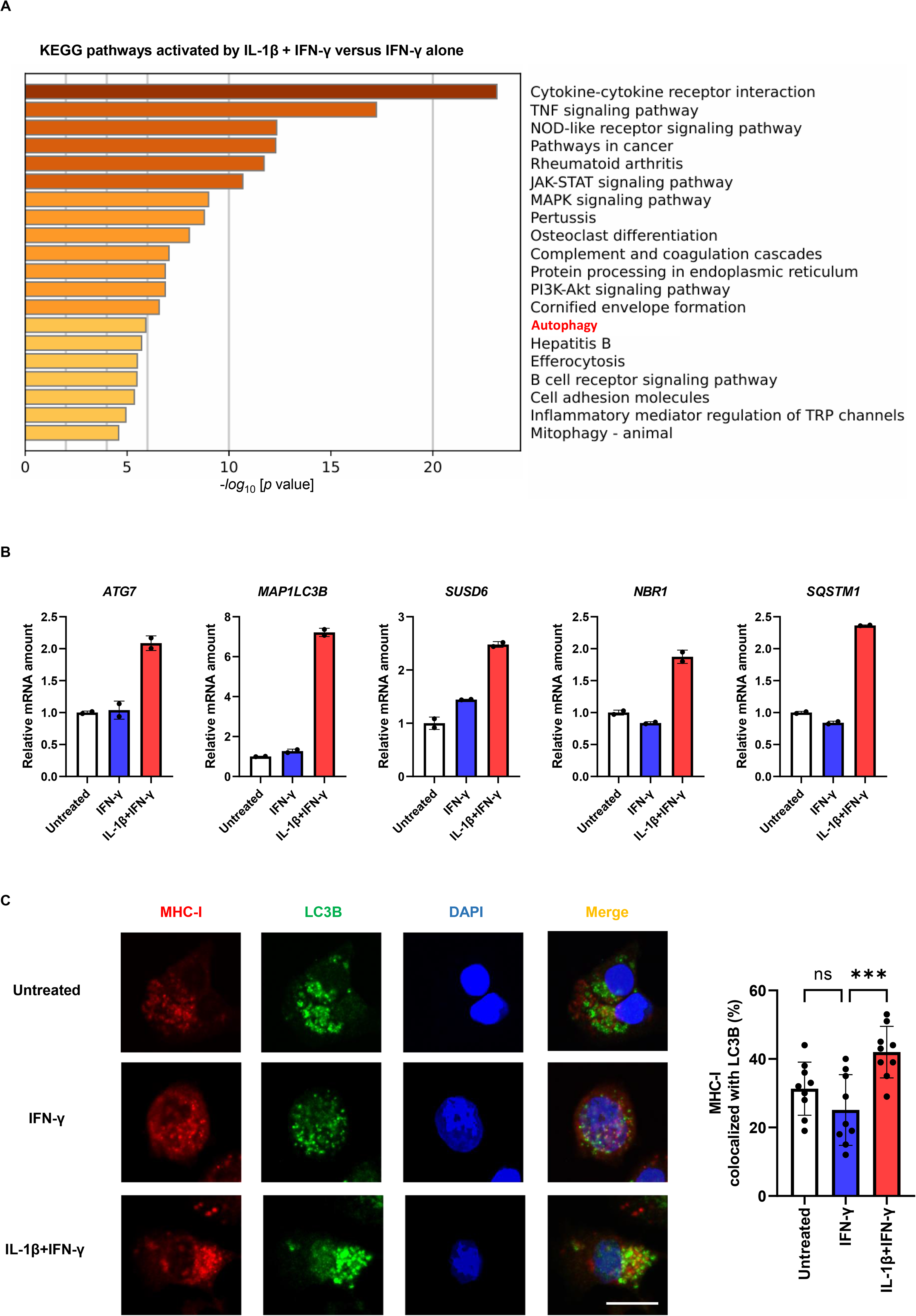
IL-1β and IFN-γ co-stimulation enhances autophagy activation and MHC-I association with autophagic structures. IL-1β and IFN-γ costimulation induces autophagy activation and MHC-I association with autophagic structures. **A:** Bulk RNA-seq analysis was performed for H1437 cells treated with IFN-γ (50 ng/ml) with or without IL-1β (50 ng/ml) or left untreated for 24 h. The top 20 pathways activated by IL-1β and IFN-γ costimulation relative to IFN-γ stimulation are shown for Kyoto Encyclopedia of Genes and Genomes (KEGG) pathway analysis. **B:** Expression of autophagosome-related genes as determined by the bulk RNA-seq analysis in **A**. Data are means ± SD from two biological replicates. **C:** H1437 cells incubated in the absence or presence of IFN-γ (50 ng/ml) and IL-1β (50 ng/ml) for 48 h and then in the additional presence of chloroquine (50 μM) for 24 h were subjected to immunofluorescence analysis of MHC-I and the autophagosome marker LC3B. Nuclei were stained with DAPI. Scale bar, 10 μm. Representative images as well as quantitation of the percentage colocalization of MHC-I with LC3B are shown. Quantitative data are means ± SD for individual cells in three biological replicates. \*\*\**p* < 0.001; ns, not significant (one-way ANOVA followed by Dunnett’s test).

### A SUSD6-SQSTM1 axis mediates MHC-I internalization in response to IL-1β plus IFN-γ

Finally, we examined whether autophagy is directly involved in the loss of surface MHC-I expression induced by IL-1β in IFN-γ–treated cells and attempted to identify MHC-I–associated molecules that might be indispensable for this process. Treatment of H1437 cells with the autophagy inhibitor 3-methyladenine did not affect surface MHC-I expression in the presence of both IL-1β and IFN-γ (Fig. 6A). Whereas our RNA-seq analysis revealed that the gene for the autophagosome marker LC3B was upregulated by IL-1β and IFN-γ costimulation (Fig. 5B), knockdown of LC3B by siRNA transfection also had no effect on surface MHC-I expression under this condition (Fig. 6B). Furthermore, treatment of H1437 cells with chloroquine, the lysosomal inhibitor, did not affect surface MHC-I expression (Fig. 6C). We performed immunoprecipitation analysis to identify potential MHC-I binding proteins that might activate the autophagy-lysosome pathway from among the autophagy-related molecules whose genes were upregulated by IL-1β and IFN-γ in our RNA-seq analysis. We found that the autophagy receptors NBR1 and SQSTM1 co-immunoprecipitated with MHC-I from H1437 cells treated with both IL-1β and IFN-γ in the presence of the lysosome inhibitor chloroquine (Fig. 6D).

**Figure 6.**
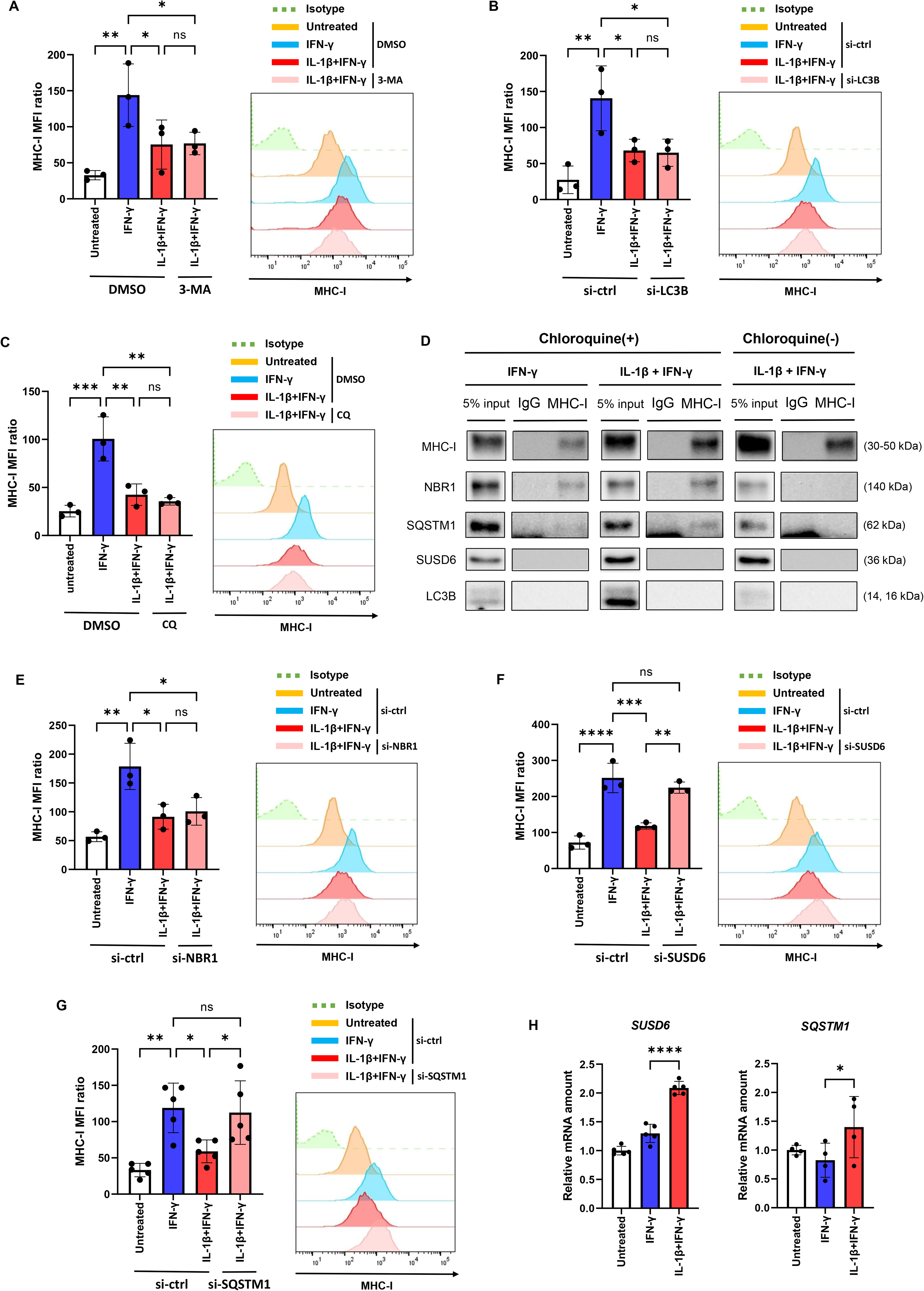
A SUSD6-SQSTM1 axis mediates MHC-I internalization in response to IL-1β plus IFN-γ. SQSTM1 associates with MHC-I and is essential for its autophagy-dependent downregulation induced by cytokine costimulation. **A, C:** Flow cytometric analysis of surface MHC-I expression in H1437 cells that had been incubated in the absence or presence of IFN-γ (50 ng/ml) and IL-1β (50 ng/ml) for 24 h and then in the additional presence of the autophagy inhibitor 3-methyladenine (3-MA, 50 mM) (A), the lysosome inhibitor chloroquine (CQ, 50 μM) (C), or DMSO vehicle for 24 h. **B, E–G:** H1437 cells were transiently transfected with LC3B (si-LC3B) (**B**), NBR1 (**E**), SUSD6 (**F**), SQSTM1 (**G**), or control (si-ctrl) siRNAs for 24 h and then treated with IFN-γ (50 ng/ml) alone or together with IL-1β (50 ng/ml) for 48 h before evaluation of surface MHC-I expression by flow cytometry. **D:** H1437 cells incubated with or without IFN-γ (50 ng/ml) and IL-1β (50 ng/ml) for 24 h and then in the additional absence or presence of chloroquine (50 μM) for 24 h were subjected to immunoprecipitation with antibodies to MHC-I or control IgG. The resulting precipitates as well as a portion of the original cell lysates (5% input) were subjected to immunoblot analysis with antibodies to the indicated proteins. Data are representative of three biological replicates. **H:** RT-qPCR analysis of SUSD6 and SQSTM1 mRNAs in H1437 cells treated for 24 h with IL-1β (50 ng/ml) or IFN-γ (50 ng/ml) as indicated All quantitative data are means ± SD from three to five independent experiments. \**p* < 0.05, \*\**p* < 0.01, \*\*\**p* < 0.001, \*\*\*\**p* < 0.0001; ns, not significant (one-way ANOVA followed by Dunnett’s test or Šídák’s multiple comparisons test).

To determine whether NBR1 and SQSTM1 are required for internalization of MHC-I, we examined the effects of their knockdown. Depletion of NBR1 in cells treated with IL-1β and IFN-γ did not restore surface MHC-I expression (Fig. 6D). However, such knockdown of SQSTM1 as well as that of SUSD6 restored surface MHC-I expression to levels similar to that apparent in cells treated with IFN-γ alone (Fig. 6E, F). Consistent with these findings and our RNA-seq data (Fig. 5B), RT-qPCR analysis revealed that costimulation with IL-1β plus IFN-γ for 24 h resulted in a significant increase in the expression of both *SUSD6* and *SQSTM1* compared with that apparent in cells treated with IFN-γ alone (Fig. 6G). Together, these observations indicated that IL-1β plus IFN-γ activate a SUSD6-SQSTM1 axis that mediates surface MHC-I downregulation via ubiquitination and degradation dependent on the autophagosome-lysosome pathway, resulting in immune evasion by NSCLC cells (Supplementary Fig. S4).

## Discussion

In the present study, we first showed that IL-1β, an inflammatory cytokine abundant within the TME of NSCLC ^22^, suppresses the IFN-γ–induced expression of MHC-I at the surface of NSCLC cells. This suppression is not mediated through inhibition of the classical mechanism of antigen presentation mediated by TAP1/2 or through inhibition of MHC-I gene transcription or protein synthesis. Rather, our immunofluorescence findings indicated that it is mediated by a posttranslational mechanism that involves increased internalization of surface MHC-I molecules. Comprehensive gene expression analysis revealed that combined IL-1β and IFN-γ stimulation selectively upregulates the expression of autophagy-related genes including *SUSD6*, which in turn promotes SQSTM1-dependent internalization of MHC-I. Our results thus uncover a tumor cell–intrinsic immune regulatory mechanism in which the inflammatory cytokine IL-1β promotes reduction of MHC-I surface expression via SQSTM1. This pathway allows cancer cells to attenuate antigen presentation, dampen immunogenicity, and may enhance cell survival under inflammatory pressure in an immune-inflamed TME with abundant IFN-γ–producing T cells such as that in NSCLC.

Autophagy is a highly conserved intracellular degradation system that plays a central role in maintenance of cellular homeostasis through the selective removal of damaged or superfluous organelles. Selective autophagy pathways—such as mitophagy, ER-phagy, and pexophagy—ensure the timely elimination of dysfunctional organelles, thereby preventing the accumulation of reactive oxygen species and protein aggregates and consequent metabolic stress. Autophagy thus supports bioenergetic balance, redox homeostasis, and cellular signaling required for the survival of various cell types ^27^ ^28^. In the context of the TME, autophagy-dependent organelle maintenance allows cancer cells to adapt to hypoxia, nutrient deprivation, and inflammatory stress ^29^. In addition, autophagy-dependent degradation of MHC-I has recently emerged as an important immune evasion pathway in cancer cells ^30^. In pancreatic and colorectal cancer, tumor cells internalize MHC-I into autophagosomes for lysosomal degradation, thereby reducing surface expression of MHC-I and impeding recognition by CD8⁺ T cells ^31^. This process is dependent on the membrane protein SUSD6, which was originally identified as a negative regulator of MHC-I at the level of ubiquitination and lysosomal trafficking in acute myeloid leukemia and solid tumors ^32^. Expression of SUSD6 was also shown to be induced by hypoxia or ER stress, resulting in ubiquitination of MHC-I, its autophagic transport via the adapter protein NBR1, and its subsequent autophagosome-lysosome–dependent degradation ^33^. These various observations revealed a regulatory mechanism governing MHC-I expression in cancer cells within an immune-desert TME such as that of pancreatic cancer or highly hypoxic tumors.

Our RNA-seq analysis showed that IL-1β and IFN-γ costimulation increased expression of canonical autophagy-related genes such as *ATG7* and *MAP1LC3B* as well as that of *SUSD6* and the adapter protein genes *NBR1* and *SQSTM1*. The expression of SQSTM1 has previously been known to be induced by a wide range of upstream stimuli, including inflammatory cytokines such as IL-1β and tumor necrosis factor–α, oxidative stress, proteasome dysfunction, ER stress, nutrient signaling and the mTORC1 (mechanistic target of rapamycin complex 1) pathway, and pathogen-associated molecular patterns ^34^ ^35^. These diverse inputs converge on transcriptional and posttranslational regulation of SQSTM1, reflecting its central role in adaptation to cellular stress. SQSTM1 functions as a selective autophagy receptor, binding to polyubiquitinated proteins and delivering them to LC3-positive autophagosomes. It also serves as a signaling scaffold that interfaces with nuclear factor–κB, mTOR, and antioxidant pathways ^36,37^. These dual functions allow SQSTM1 to integrate inflammatory, metabolic, and proteotoxic stress signals in order to promote cell survival and homeostasis.

Our immunoprecipitation data revealed that, in the presence of the lysosome inhibitor chloroquine, MHC-I was associated with NBR1 and SQSTM1 in NSCLC cells. Furthermore, we found that knockdown of SUSD6 or SQSTM1, but not that of NBR1, or autophagy inhibitor 3-MA, knockdown of LC3B, lysosomal inhibitor chloroquine, restored MHC-I surface expression in costimulated cells to levels similar to that apparent in cells treated with IFN-γ alone. These observations suggest that SQSTM1 plays a non-redundant role in the early handling of MHC-I cargo, potentially at a pre-autophagic stage ^36,37^. One possible interpretation is that, under inflammatory cytokine costimulation in immune-inflamed tumors, MHC-I molecules undergo SUSD6-mediated ubiquitination and are subsequently recognized by SQSTM1, which facilitates their sequestration into condensate-like structures and promotes internalization. In this context, SQSTM1-dependent cargo condensation may represent a critical step in reducing surface MHC-I availability, independently of, or prior to, downstream autophagosome–lysosome–mediated degradation. Consistent with this, SQSTM1 has been shown to organize ubiquitinated proteins into higher-order assemblies that serve as substrates for subsequent selective autophagy^36^ ^37^. Our findings suggest that the stress-adaptive machinery dependent on SQSTM1 is co-opted by cancer cells. Under conditions in which T cells produce IFN-γ in order to eliminate tumor cells and in which immunosuppressive cells such as TAMs produce IL-1β, SQSTM1-dependent internalization of MHC-I and attenuation of surface MHC-I expression in cancer cells occur. In this context, SQSTM1 may function as a molecular hub that links stress tolerance to immune escape in an immune-inflamed TME.

Several questions remain with regard to the immunoregulatory pathway identified in our study. First, we have not evaluated whether the IL-1β–SUSD6–SQSTM1–MHC-I axis influences CD8⁺ T cell infiltration, antigen recognition, or cytotoxic activity. Future comprehensive studies incorporating protein-level, cell-surface, and spatial readouts—such as MHC-I surface expression, SQSTM1 localization, and cytokine profiling in patient samples—will be required to determine whether this pathway operates in vivo. Second, clarification of upstream signaling mechanisms governing IL-1β–induced SUSD6 expression and the molecular criteria for MHC-I substrate recognition is required. Third, the downstream fate of internalized MHC-I combined with SQSTM1 or the execution mechanism of autophagy-lysosome pathways warrants further investigation. Finally, characterization of how other major inflammatory cytokines in the TME, such as IL-6 and tumor necrosis factor–α, might influence surface expression of MHC-I in cancer cells will be essential for defining the broader cytokine architecture governing tumor immune evasion.

## Conclusion

By studying cell models of immune-inflamed lung cancer, we have shown that IL-1β counteracts IFN-γ–induced surface expression of MHC-I by promoting SQSTM1-dependent internalization of MHC-I, revealing a previously underappreciated mechanism of tumor cell–intrinsic immune regulation under inflammatory conditions. Future studies aimed at delineating the downstream molecular machinery of SQSTM1, including its potential coupling to ubiquitin-dependent pathways and in vivo immune contexts, will be essential to fully define the impact of this axis on tumor immunogenicity.

## Abbreviations

ANOVA: analysis of variance
CHX: cycloheximide
CTL: cytotoxic T lymphocyte
DAPI: 4΄,6-diamidino-2-phenylindole
DMSO: dimethyl sulfoxide
ER: endoplasmic reticulum
IFN-γ: interferon-γ
IgG: immunoglobulin G
IL-1β: interleukin-1β
MFI: mean fluorescence intensity
MHC-I: major histocompatibility complex class I
NSCLC: non–small cell lung cancer
PBS: phosphate-buffered saline
PD-L1: programed cell death–ligand 1
qPCR: quantitative polymerase chain reaction
RT: reverse transcription
seq: sequencing
siRNA: small interfering RNA
TAM: tumor-associated macrophage
TME: tumor microenvironment

## Declarations

### Ethics approval and consent to participate

Not applicable.

### Consent for publication

Not applicable.

### Availability of data and materials

The datasets analyzed during the current study are available from the corresponding author on reasonable request.

### Authors’ contributions

SY and K. Tanaka conceived and designed the study as well as contributed to writing of the manuscript. SY, HS and AH performed all experiments with the assistance of K. Tanaka and K. Tsubouchi. SY, K. Tanaka, and AH analyzed the data. All authors contributed to data interpretation and revision of the article and approved the submitted version.

## Acknowledgements

We thank staff of Cell Innovator (Fukuoka, Japan) for helpful discussion and technical support.

---

**Supplementary Fig. S1.**
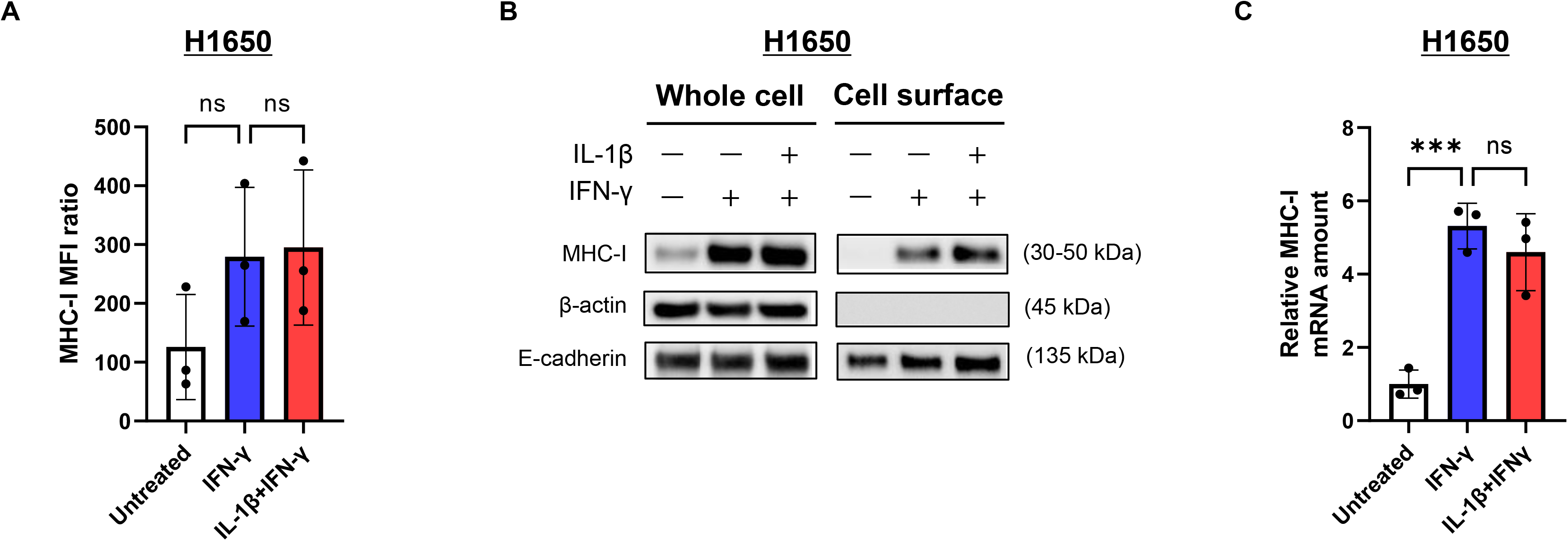
MHC-I expression enhanced by IFN-γ stimulation is not suppressed by combined IL-1β stimulation in H1650 cells. IFN-γ–induced surface MHC-I expression is not suppressed by IL-1β in H1650 cells. **A:** Flow cytometric analysis of MHC-I expression at the cell surface in H1650 cells treated with IFN-γ (50 ng/ml) either alone or together with IL-1β (50 ng/ml) for 48 h. **B:** Immunoblot analysis of MHC-I in whole cell lysates and a cell surface protein extract prepared from H1650 cells treated as in **A**. Data are representative of three biological replicates. **C:** RT-qPCR analysis of MHC-I mRNA in H1650 cells treated as in **A**. Quantitative data in **A** and **C** are means ± SD from three independent experiments. \*\*\**p* < 0.001; ns, not significant (one-way ANOVA followed by Dunnett’s test).

**Supplementary Fig. S2.**
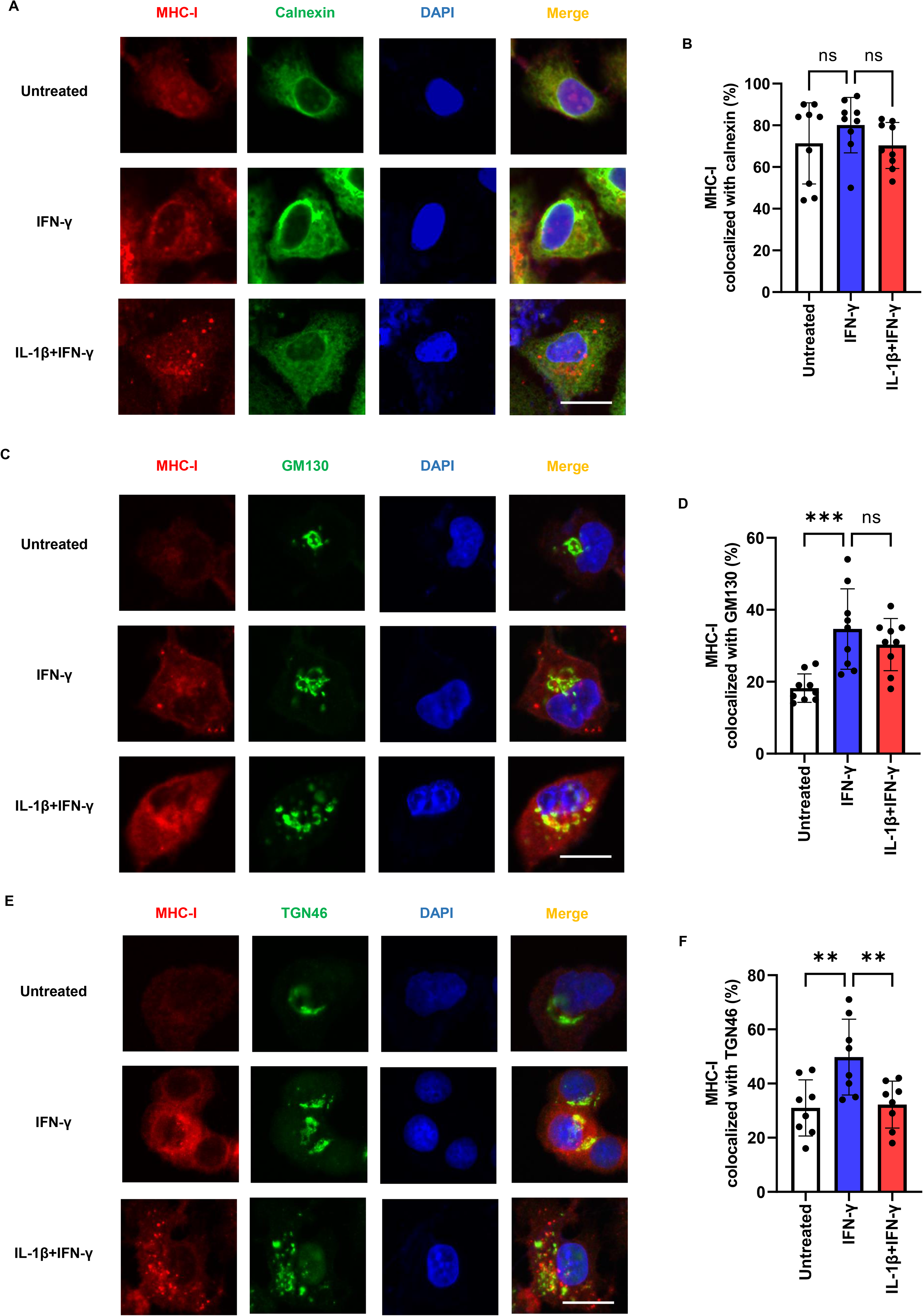
MHC-I exocytosis was maintained under combined IL-1β stimulation. MHC-I transport is maintained in the presence of IL-1β. H1437 cells incubated in the absence or presence of IFN-γ (50 ng/ml) and IL-1β (50 ng/ml) for 48 h were fixed, permeabilized, and subjected to immunofluorescence analysis of MHC-I and either the ER marker calnexin (**A**), the cis-Golgi marker GM130 (**C**), or the trans-Golgi marker TGN46 (**E**). Nuclei were stained with DAPI. Scale bars, 10 µm. Quantification of the percentage colocalization of MHC-I with the three markers is shown in **B**, **D**, and **F**, respectively. The quantitative data are means ± SD for individual cells from three independent experiments. \*\**p* < 0.01, \*\*\**p* < 0.001; ns, not significant (one-way ANOVA followed by Dunnett’s test).

**Supplementary Fig. S3.**
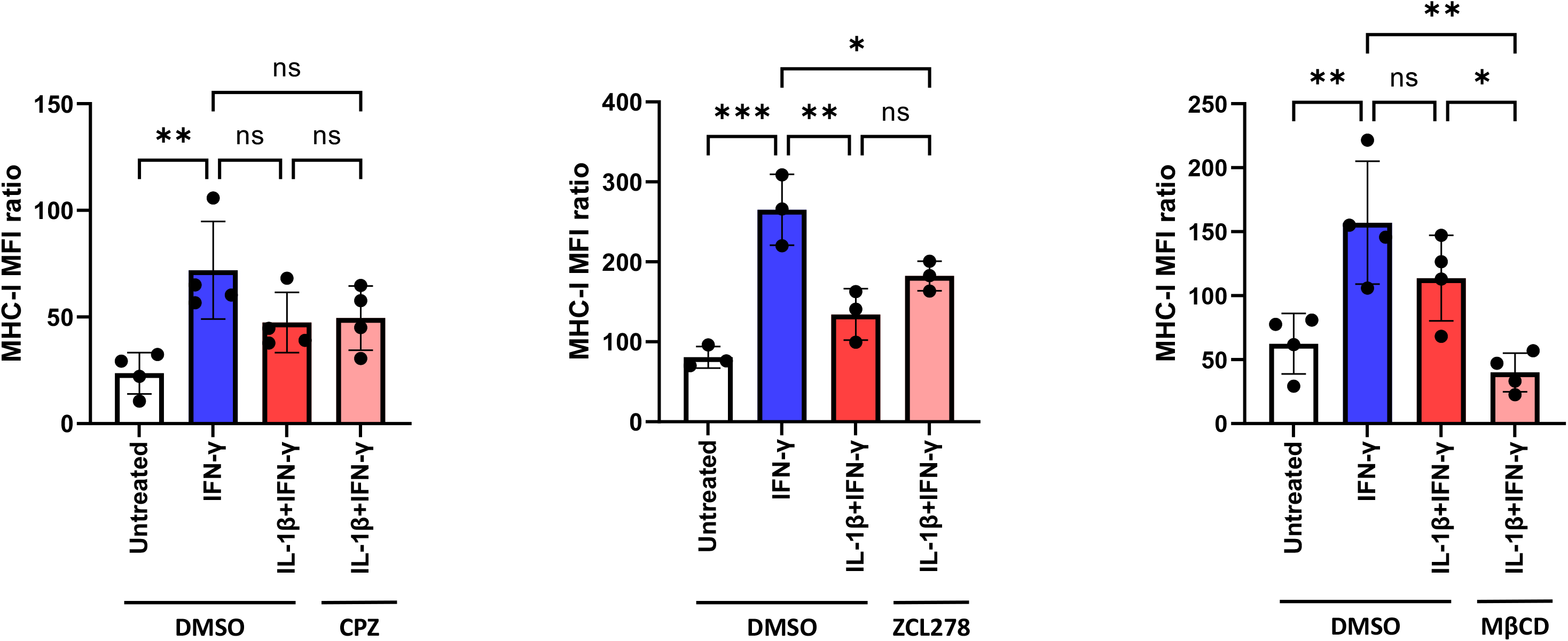
Endocytosis did not appear to be involved in the suppression of MHC-I expression by IL-1β co-stimulation. Suppression of surface MHC-I expression by IL-1β is independent of endocytosis pathways. H1437 cells were incubated in the absence or presence of IFN-γ (50 ng/ml) and IL-1β (50 ng/ml) for 24 h and then in the additional absence or presence of chlorpromazine (CPZ, 50 μM), ZCL278 (10 μM), methyl-β-cyclodextrin (MβCD, 5 mM), or DMSO vehicle for 3 h. Surface MHC-I expression was then quantified by flow cytometry. Data are means ± SD from three or four independent experiments. \**p* < 0.05, \*\**p* < 0.01, \*\*\**p* < 0.001; ns, not significant (one-way ANOVA followed by Dunnett’s test).

**Supplementary Fig. S4.**
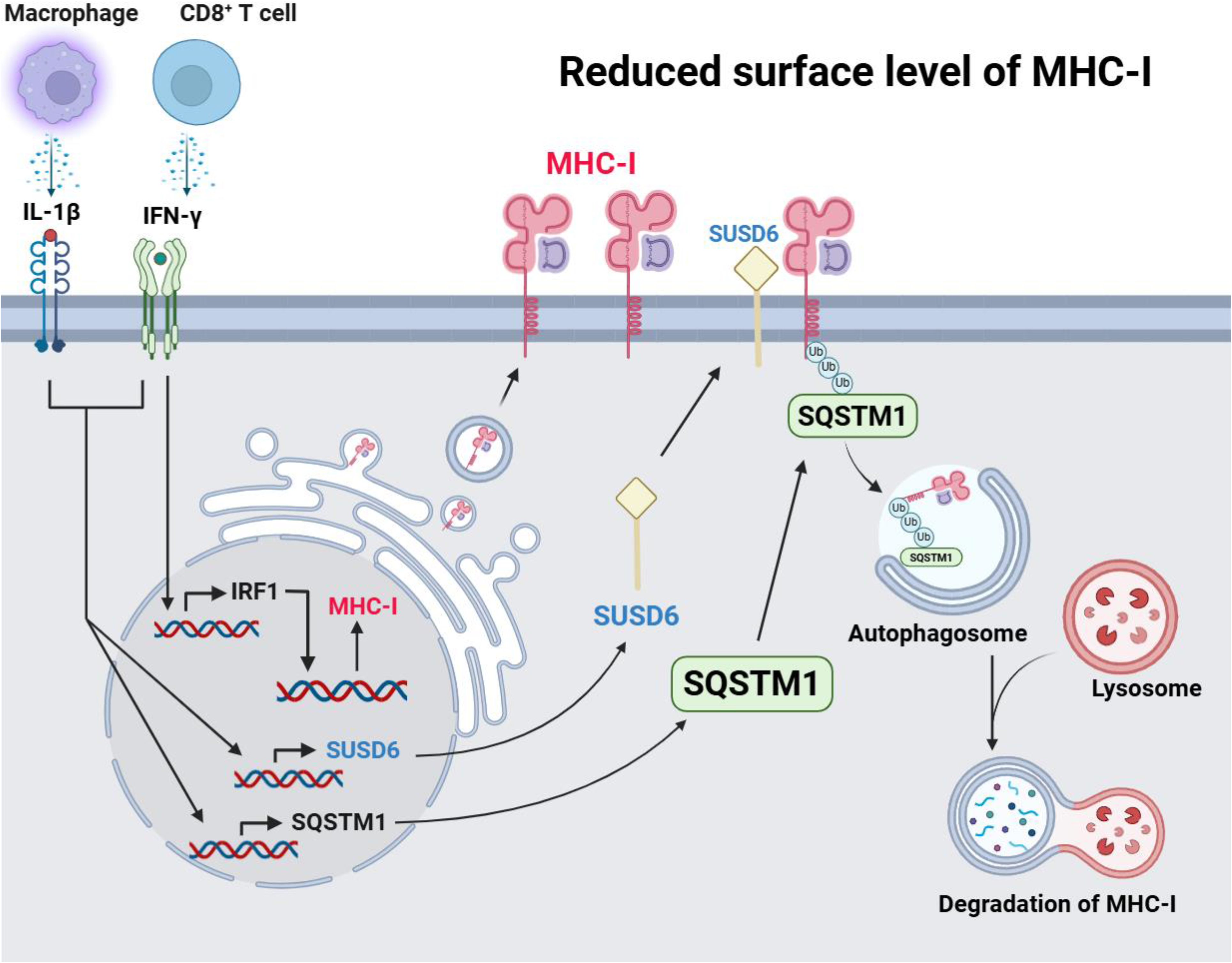
Graphical summary showing the mechanism of cytokine-driven SUSD6/SQSTM1 axis in the reduction of MHC-I surface expression. Proposed model for the role of a SUSD6-SQSTM1 axis in the downregulation of surface MHC-I expression by IL-1β in IFN-γ–stimulated NSCLC cells. IFN-γ derived from T cells and IL-1β derived from macrophages together induce transcription of *SUSD6* and *SQSTM1* in tumor cells, resulting in downregulation of MHC-I at the cell surface via SUSD6-dependent endocytosis and its SQSTM1-dependent degradation via the autophagy-lysosome pathway.

